## Supplemental Information for "Dissecting the unique self-assembly landscape of the HIV-2 capsid protein"

**Table S1.** Protein constructs purified in this study. All constructs were expressed using pET-11a (ampicillin resistant) bacterial expression vectors.

| Construct | Purpose |
|---|---|
| HIV-2 GL-AN CA | WT HIV-2 CA for assembly and morphological comparison |
| HIV-2 GL-AN CA P14C/E45C | Disulfide stabilization of assembled HIV-2 CA lattice |
| HIV-2 GL-AN CA P14C/E45C/W184A/M185A | Disulfide stabilization of HIV-2 CA hexamers |
| HIV-2 GL-AN CA-TEV-MBP-6xHis L26V/G39M | Assembly and morphological comparison of an HIV-1-mimicking mutation at the NTD-NTD interface. Tags for solubility. |
| HIV-2 GL-AN CA K31A | Assembly and morphological comparison of an HIV-1-mimicking mutation at the NTD-CTD interface. |
| HIV-2 GL-AN CA Q41S | Assembly and morphological comparison of an HIV-1-mimicking mutation at the NTD-NTD interface. |
| HIV-2 GL-AN CA Q41E/N57D | Morphological comparison of a pentamer-disrupting mutation. |
| HIV-2 GL-AN CA A42C/Q54C | Disulfide stabilization of assembled HIV-2 CA lattice |
| HIV-2 GL-AN CA Y50Q | Assembly and morphological comparison of an HIV-1-mimicking mutation at the NTD-NTD interface and morphological comparison of a hexamer-disrupting mutation. |
| HIV-2 GL-AN CA Y50Q/K170N | Morphological comparison of a hexamer-disrupting mutation. |
| HIV-2 GL-AN CA D61G | Assembly and morphological comparison of an HIV-1-mimicking mutation at the NTD-CTD interface. |
| HIV-2 GL-AN CA K170A | Morphological comparison of a hexamer-disrupting mutation. |
| HIV-2 GL-AN CA K170N | Morphological comparison of a hexamer-disrupting mutation. |
| HIV-2 GL-AN CA D178S/P179Q/A180E | Assembly and morphological comparison of an HIV-1-mimicking mutation at the CTD-CTD interface. |
| HIV-1 NL4-3 CA | WT HIV-1 CA for assembly and morphological comparison |
| HIV-1 NL4-3 CA-TEV-MBP-6xHis A14C/E45C | Disulfide stabilization of assembled HIV-1 CA lattice. Tags for solubility. |

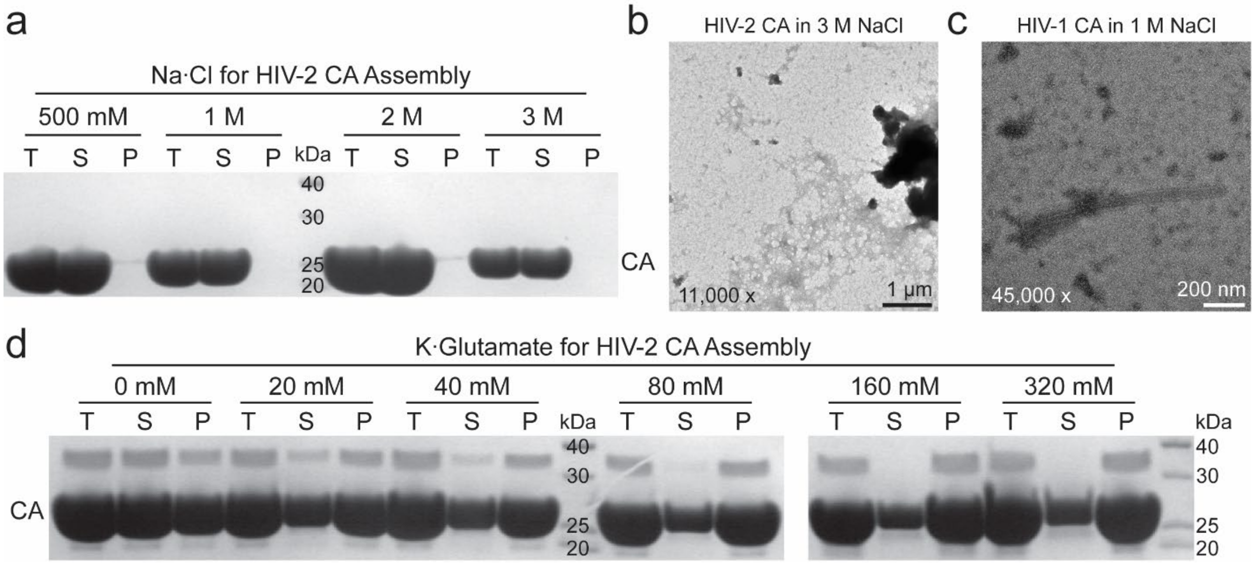


**Figure S1.** Evidence of in vitro assembly of WT HIV-2 capsid protein (CA). (a) Sedimentation assay of WT HIV-2 CA assembled at high protein (500 μM) and inositol hexakisphosphate (IP6) (2.5 mM) concentrations, with varying concentrations of NaCl. Abbreviations are as follows: T - Total; S - Supernatant; P - Pellet. (b) Representative negative-stain electron microscopy (EM) micrograph of WT HIV-2 CA assembled in 3 M NaCl, showing no evidence of ordered assemblies. (c) Representative negative-stain EM micrograph of WT HIV-1 CA assembled in 1 M NaCl, demonstrating the formation of tubular structures. (d) Sedimentation assay of WT HIV-2 CA assembled at high protein (500 μM) and IP6 (2.5 mM) concentrations with varying potassium glutamate (KGlu) concentrations, similar to (a). (e) Representative cryo-EM micrograph of WT HIV-2 CA (500 μM) assembled in the presence of 2.5 mM IP6, and 1 M KGlu.

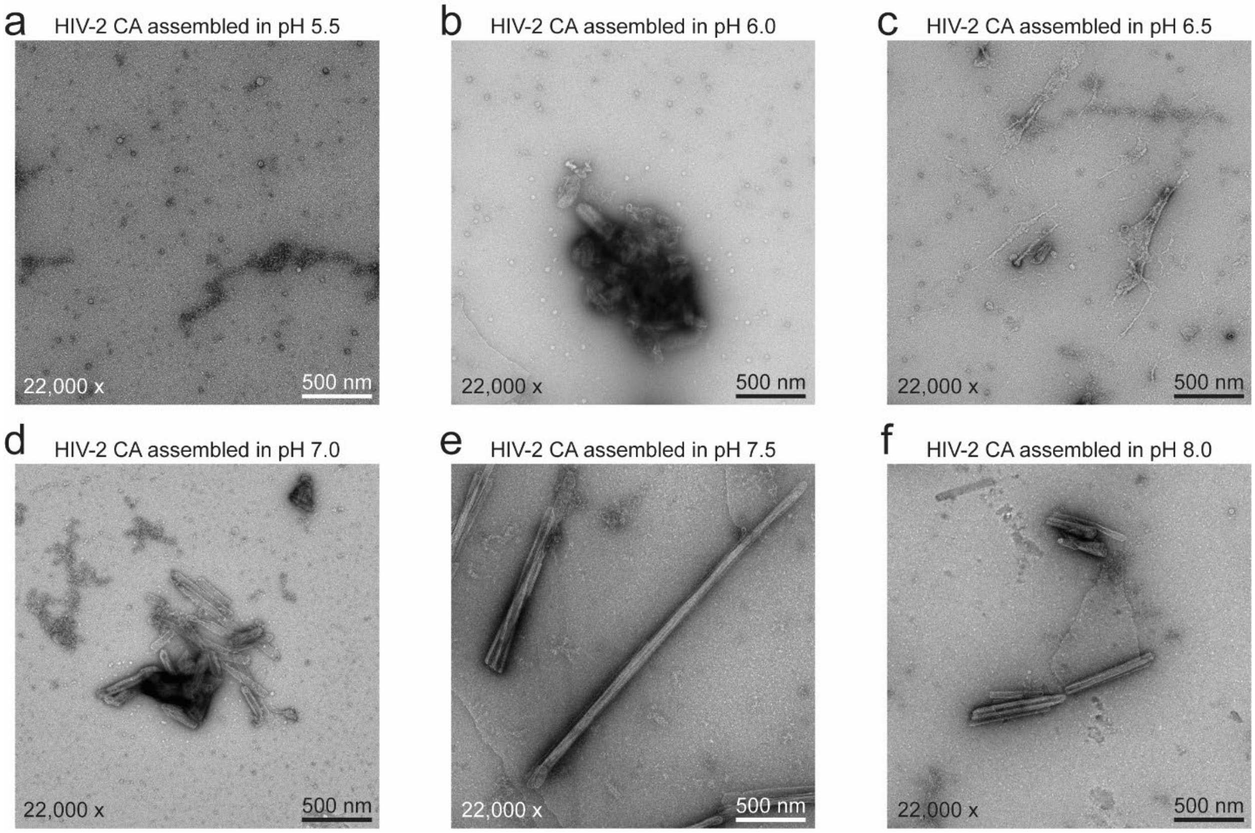


**Figure S2.** HIV-2 CA assembly morphology as a function of buffer pH. Representative negative-stain EM micrographs of WT HIV-2 CA assembled with 1 M KGlu under the following buffer pH conditions: (a) pH 5.5; (b) pH 6.0; (c) pH 6.5; (d) pH 6.5; (e) pH 7.0; (f) pH 7.5; (g) pH 8.0.

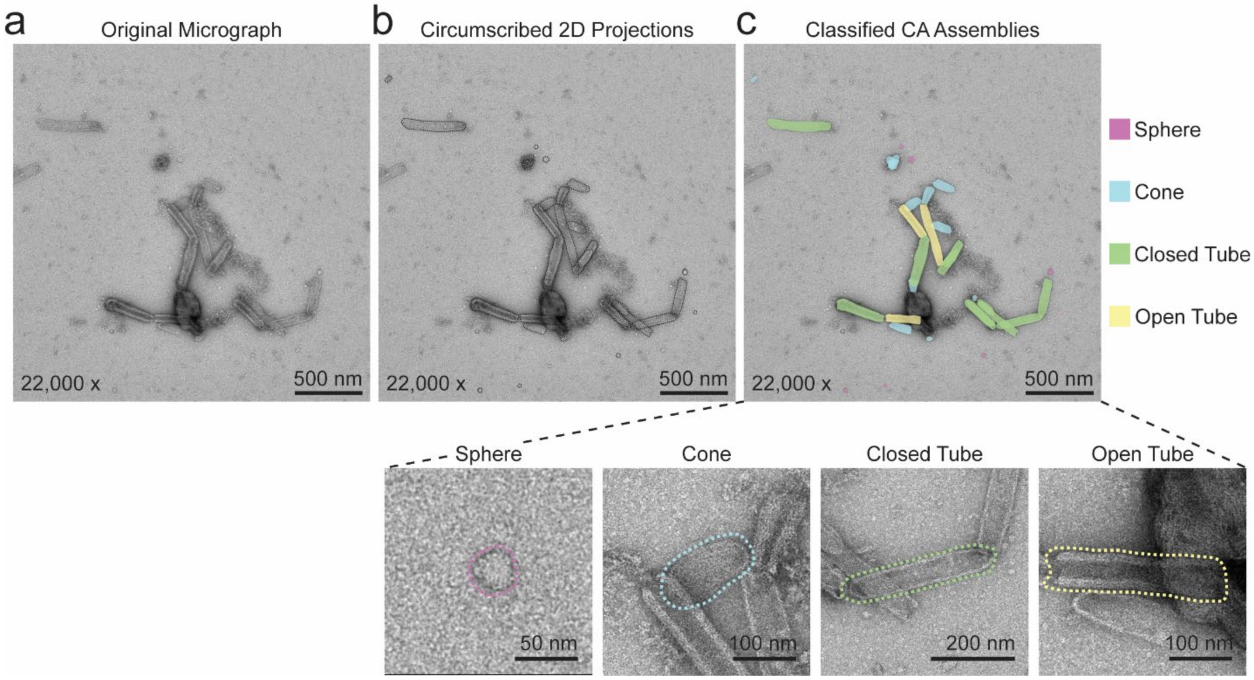


**Figure S3.** Strategies for quantifying CLP morphology. Example negative-stain EM micrograph of assembled WT HIV-2 to illustrate the morphology quantification workflow. (a) Unmodified micrograph. (b) Particle outlines overlaid on the original micrograph, illustrating the method used to circumscribe 2D particle projections for area calculations. (c) Shaded areas superimposed on the original micrograph to demonstrate how individual particles would be classified for morphology comparisons.

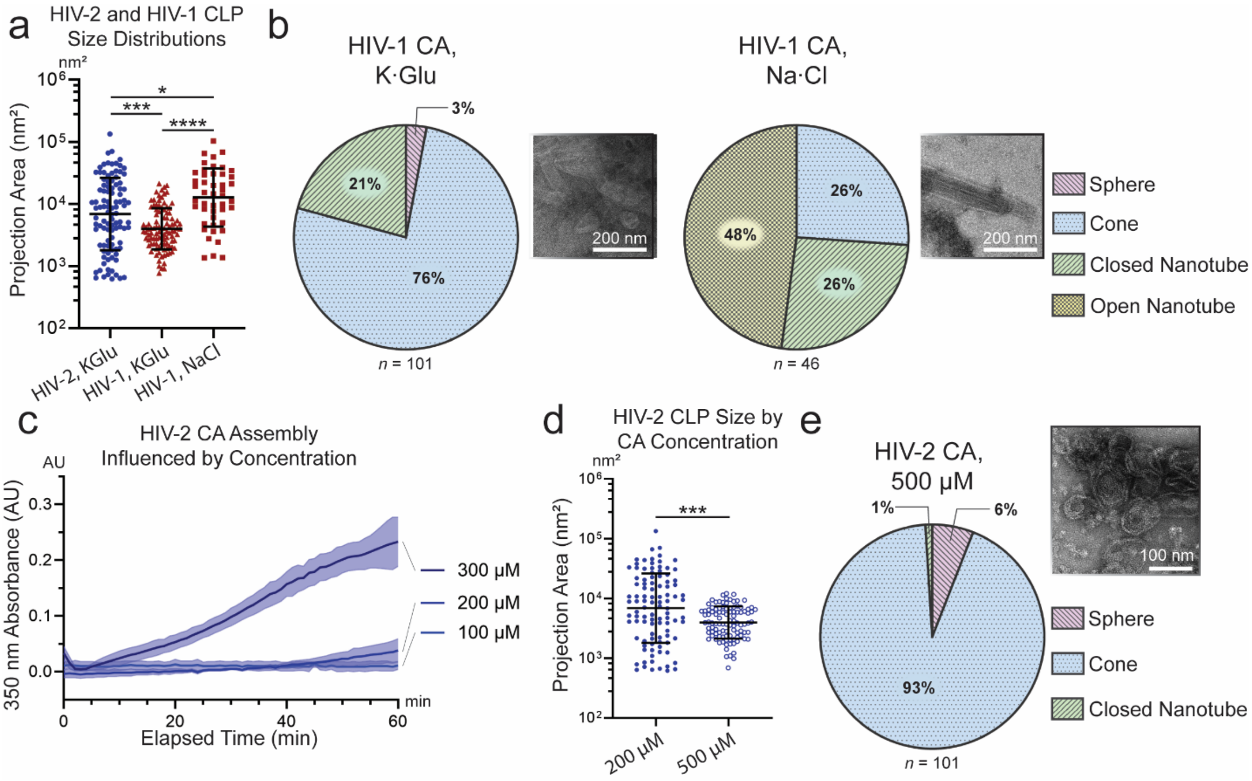


**Figure S4.** Differential effects of salt and concentration on in vitro assembly of HIV CAs. (a) Sizes of in vitro assemblies of WT HIV-1 CA induced by varying salt conditions, measured by 2D projection area on a logarithmic axis. The central line indicates the geometric mean, with the upper and lower lines representing geometric standard deviations from the mean (Significance values: * = $p < 0.05$; *** = $p < 0.001$; **** = $p < 0.0001$). (b) Proportions of in vitro assemblies of HIV-1 CA induced by different salts, classified by particle morphology. Representative fragments of micrographs are shown to the right of the corresponding chart. Full micrographs are provided in Figure 1c (KGlu) and Figure S1c (NaCl). (c) Assembly of HIV-2 CA induced by 600 mM KGlu and monitored by absorbance at 350 nm across varying protein concentrations. The line connects mean values of adjacent time points ($n$ = 3), and the shaded area denotes standard deviation from the mean. (d) Sizes of in vitro assemblies of WT HIV-2 CA varying initial protein concentrations, measured by 2D projection area on a logarithmic axis. Data point annotations and significance values are as described in (a). (e) Proportions of in vitro assemblies of HIV-2 CA at 500 μM initial protein concentration, classified by particle classification. Upper right: representative negative-stain EM micrograph of this assembly reaction.

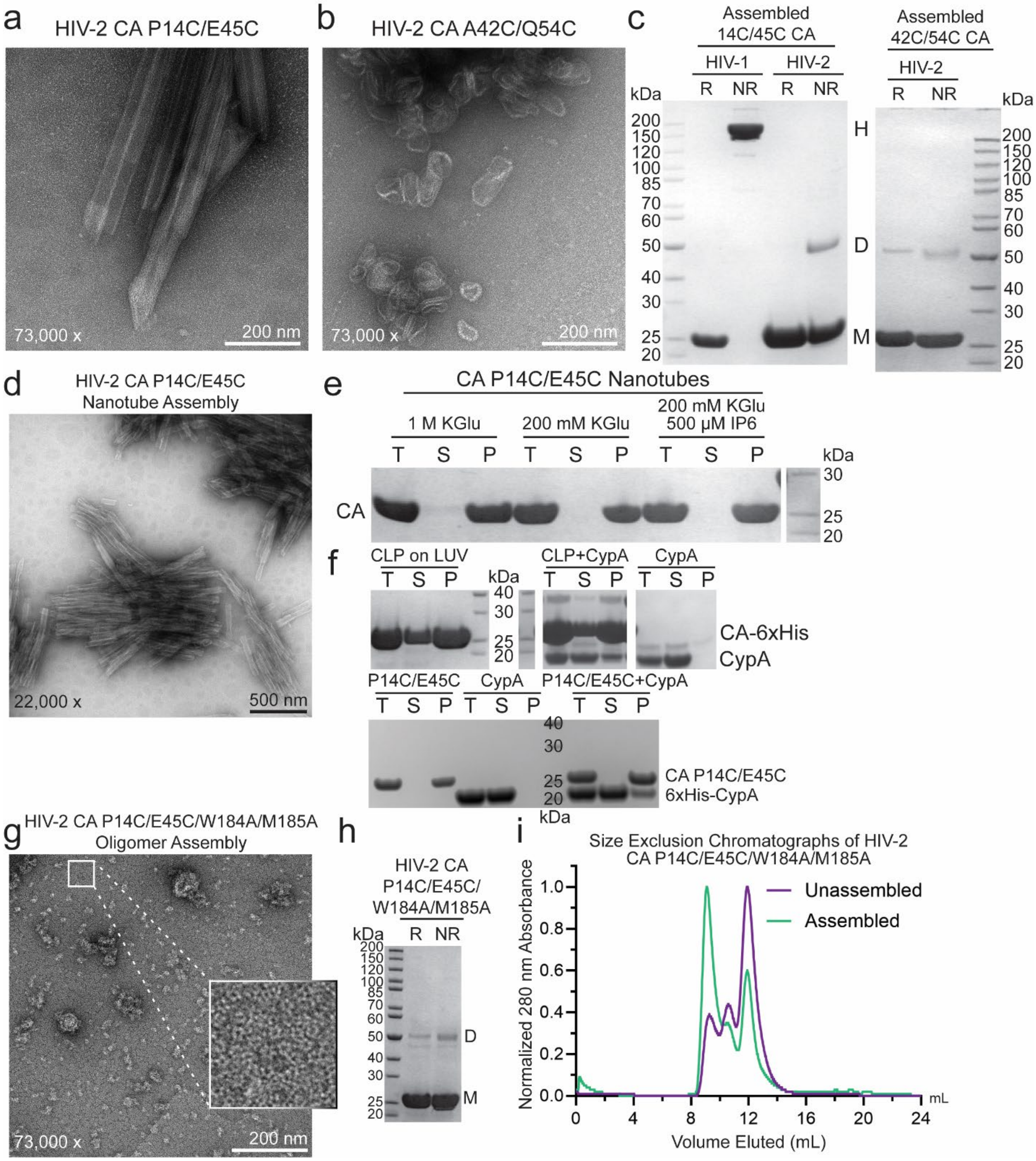


**Figure S5.** Assembly and sedimentation of HIV-2 CA nanotubes. Representative negative-stain EM micrographs of (a) assembled HIV-2 CA P14C/E45C and (b) assembled HIV-2 CA A42C/Q54C. (c) Comparison of disulfide bond formation efficiency in HIV-1 CA A14C/E45C, HIV-2 CA P14C/E45C, and HIV-2 CA A42C/Q54C, assessed by SDS-PAGE. R - reducing condition; NR - non-reducing condition. M - monomeric CA; D - dimeric CA; H - hexameric CA. (d) Representative negative-stain EM micrograph of HIV-2 CA P14C/E45C nanotubes. (e) SDS-PAGE analysis of sedimentation assay samples from assembled HIV-2 CA P14C/E45C nanotubes following dialysis into buffers containing the indicated KGlu concentrations. Abbreviations are as follows: T - Total; S - Supernatant; P - Pellet. Gel spliced to remove irrelevant lanes. (f) SDS-PAGE gels of co-sedimentation assays involving either HIV-2 CLPs templated on large unilamellar vesicles (LUVs) (above) or HIV-2 CA P14C/E45C nanotubes (below) co-sedimenting cyclophilin A (CypA), demonstrating improved pelleting efficiency of the stabilized nanotubes. Abbreviations as defined in (e). Right gel spliced to remove irrelevant lanes. (g) Representative negative-stain EM micrograph of assembled HIV-2 CA P14C/E45C/W184A/M185A, revealing apparent hexameric assemblies. (h) SDS-PAGE gel of post-assembly HIV-2 CA P14C/E45C/W184A/M185A, demonstrating poor formation of inter-CA disulfide bonds. Abbreviations as described in (c). Gel spliced to remove irrelevant lanes. (i) Size exclusion chromatography trace of HIV-2 CA P14C/E45C/W184A/M185A before and after in vitro assembly, demonstrating an increased proportion of higher molecular weight species following assembly. Absorbance values were normalized to the maximum value of each individual trace.

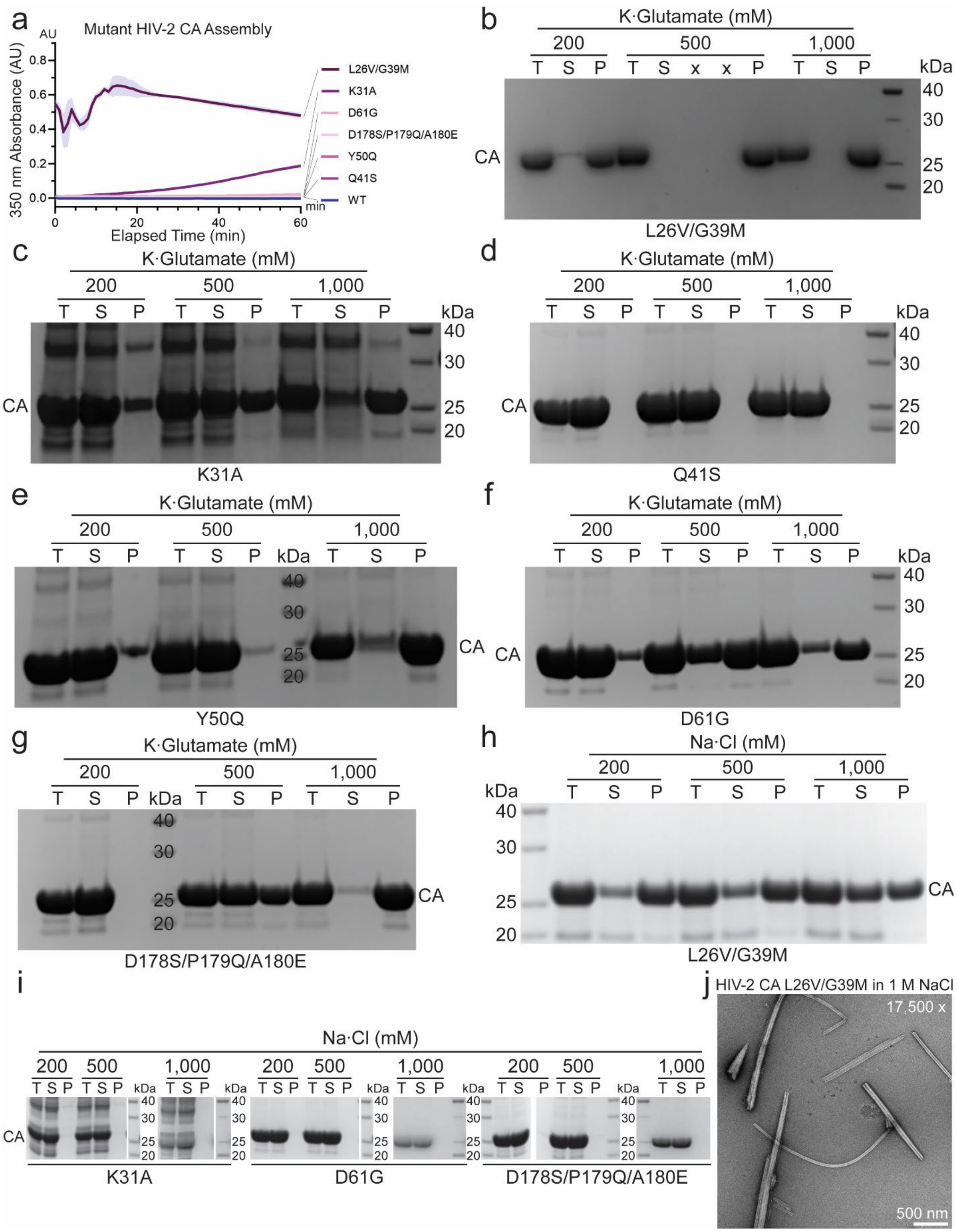


**Figure S6.** Assembly and sedimentation of HIV-2 CA mutants. (a) Assembly of WT HIV-2 CA and HIV-1-mimicking mutants induced by 500 mM KGlu and monitored by absorbance at 350 nm across varying protein concentrations. The line connects the mean values of the adjacent time points ($n$ = 3). Shaded area denotes standard deviation from the mean. Under these conditions only the highly assembly-prone mutants assembled efficiently. SDS-PAGE analysis of the sedimentation assays of the following HIV-1-mimicking HIV-2 CA mutants assembled in given KGlu concentrations: (b) L26V/G39M; (c) K31A; (d) Q41S; (e) Y50Q; (f) D61G; (g) D178S/P179Q/A180E. Abbreviations are as follows: T – Total; S – Supernatant; P – Pellet; x – no sample. (h) SDS-PAGE analysis of sedimentation assays of HIV-2 CA L26V/G39M assembled in the presence of given NaCl concentrations. (i) SDS-PAGE analysis of sedimentation assays of other HIV-2 CA mutants with improved assembly efficiency in KGlu, attempting to assemble in the presence of given NaCl concentrations. Gels spliced as needed to remove irrelevant lanes. (j) Representative negative-stain EM micrograph of CA L26V/G39M assembled in the presence of 1 M NaCl, revealing more tubular assemblies.

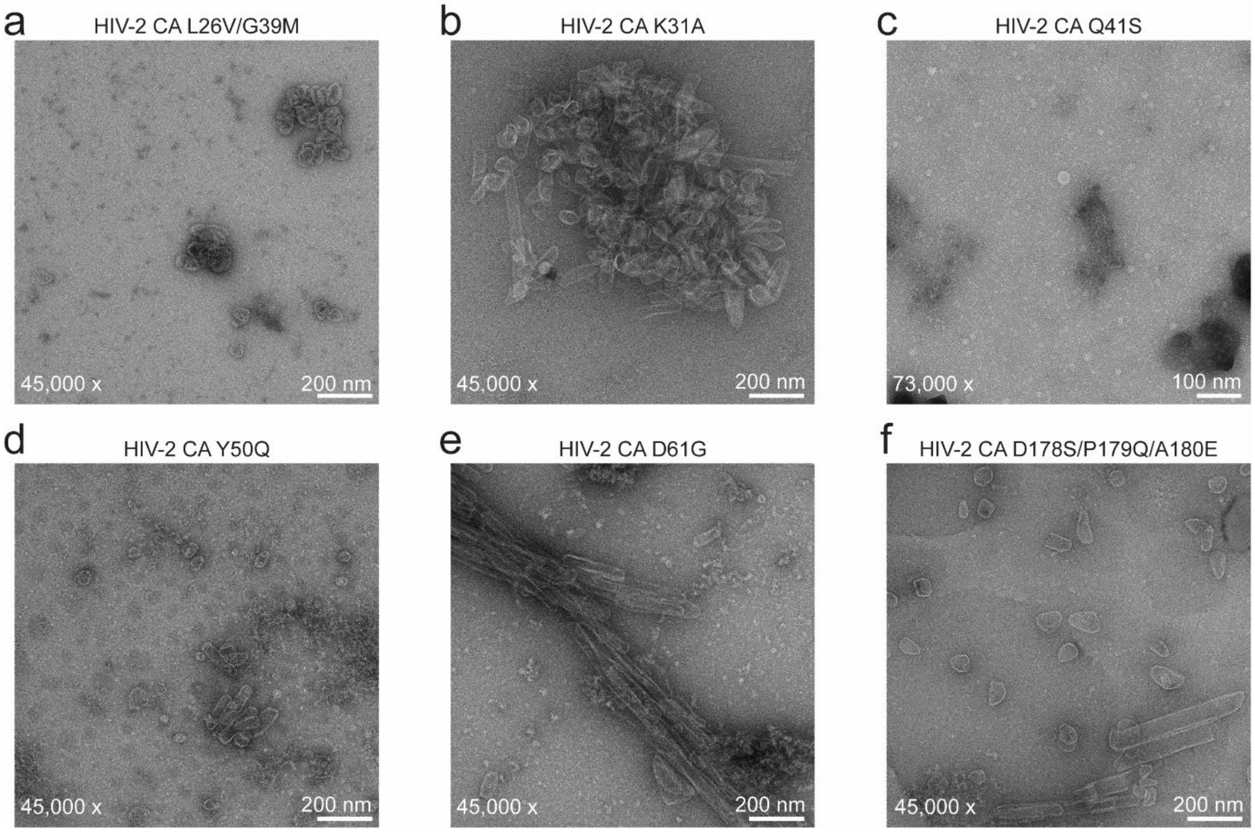


**Figure S7.** Negative-stain EM micrographs of assembled HIV-2 CA mutants. Representative negative-stain EM micrographs of assemblies of HIV-1-mimicking HIV-2 CA interface mutants: (a) L26V/G39M; (b) K31A; (c) Q41S; (d) Y50Q; (e) D61G; (f) D178S/P179Q/A180E.

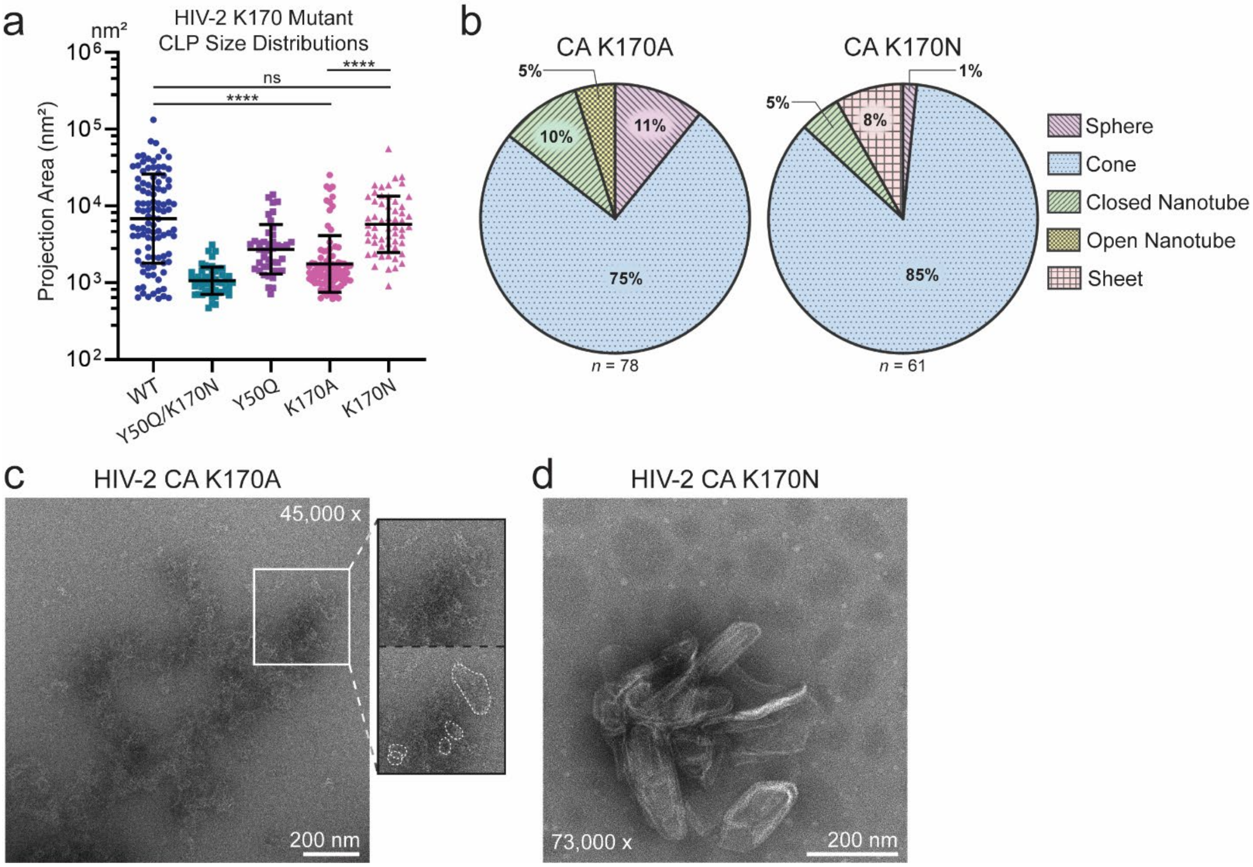


**Figure S8.** Assembly effects of HIV-2 CA K170 mutants. (a) Quantification of the areas of the 2D projections of in vitro assemblies of HIV-2 CA K170 mutants on a logarithmic axis. The central line indicates the geometric mean with upper and lower lines marking the geometric standard deviations from the mean (Significance values: ns = Not significantly different, $p > 0.05$; **** = $p < 0.0001$). (b) Proportions of in vitro assemblies of HIV-2 CA K170A (left) or K170N (right), classified by particle morphology. Both mutants predominantly formed conical assemblies. (c) Representative negative-stain EM micrograph of HIV-2 CA K170A; assembly quality appears reduced compared to other HIV-2 CA mutants. Inset highlights some assemblies amid the aggregate mass. (d) Representative negative-stain EM micrograph of HIV-2 CA K170N.
